# Binding of regulatory proteins to nucleosomes is modulated by dynamic histone tails

**DOI:** 10.1101/2020.10.30.360990

**Authors:** Yunhui Peng, Shuxiang Li, Alexey Onufriev, David Landsman, Anna R. Panchenko

## Abstract

Despite histone tails’ critical roles in epigenetic regulation, little is known about mechanisms of how histone tails modulate the nucleosomal DNA solvent accessibility and recognition of nucleosomes by other macromolecules. Here we generate extensive atomic level conformational ensembles of histone tails in the context of the full human nucleosome, totaling 26 microseconds of molecular dynamics simulations. We explore the histone tail binding with the nucleosomal and linker DNA and observe rapid conformational transitions between bound and unbound states allowing us to estimate kinetic and thermodynamic properties of the histone tail-DNA interactions. Different histone types exhibit distinct, although conformationally heterogeneous, binding modes and each histone type occludes specific DNA regions from the solvent. Using a comprehensive set of experimental data on nucleosome structural complexes, we find that majority of the studied nucleosome-binding proteins and histone tails target mutually exclusive regions on nucleosomal or linker DNA around the super-helical locations ±1, ±2, and ±7. This finding is explained within the generalized competitive binding and tail displacement models of partners recruitment to nucleosomes. Finally, we demonstrate the crosstalk between different histone post-translational modifications, where charge-altering modifications and mutations typically suppress tail-DNA interactions and enhance histone tail dynamics.

## Introduction

In eukaryotic cells, DNA is packaged in the form of chromatin and should be dynamically accessed during transcription and replication processes with high spatiotemporal precision. These seemingly contradictory tasks of DNA packaging and DNA access have been of tremendous research interest. Nucleosomes represent the basic subunits of chromatin structure and comprise a histone octamer of four types of core histones, two copies each (H2A, H2B, H3, and H4) and ~147 bp of DNA wrapped around them [1]. Intrinsically disordered histone tails flanking histone core domains play particularly important roles, and experiments show that deletions of histone tails may result in the transient unwrapping of DNA, an increase in the nucleosome sliding rate, and a decrease in nucleosome stability [2–4]. Moreover, histone tails may contribute to the inter-nucleosomal interactions and affect the higher-order chromatin structure [5–7].

Histone tails have a high degree of conformational flexibility and might protrude into the solvent and remain perpetually accessible for binding by chromatin factors [1, 8–10]. However, there is growing evidence that histone tails can extensively interact with the nucleosomal and linker DNA [11–17], which raises the possibility that tails may modulate the nucleosomal and linker DNA accessibility and regulate the nucleosome recognition by binding partners. It has been shown that despite the lower net negative charge of the nucleosome compared to the free DNA, nucleosomes are characterized by an enhanced negative charge density (so-called electrostatic focusing) even within the intact positively charged histone tails [18]. However, there are very few studies systematically characterizing the histone tail conformational ensemble in the context of the full nucleosome, physicochemical properties of their binding to DNA, and functional roles in regulatory mechanisms [11, 12, 19]. This is explained by the difficulty in experimentally observing and simulating the intrinsically disordered tails’ conformational space in the context of the full nucleosome.

Here we explore a spectrum of conformational states of disordered tails in the context of the full nucleosome to understand how conformational dynamics of histone tails modulate the DNA solvent accessibility and the recognition of nucleosome-binding partners. We perform extensive sampling of tail conformations with the native human DNA sequence totaling in 26 microseconds simulated trajectories. We find rapid interconversions between histone tail-DNA bound and unbound states and show that the ensemble of tail conformations adheres to the nucleosome two-fold symmetry requirement and provides reasonable estimates of tail-DNA dissociation constants. Finally, we utilize experimental data on nucleosome structural complexes and dissociation constants of chromatin factors binding to explore how tail dynamics may mediate or inhibit the interactions of nucleosomes with their binding partners and how tails’ post-translational modifications (PTMs) and mutations may influence this process. We find that many nucleosome-binding proteins and histone tails target overlapping and mutually exclusive regions on nucleosomal or linker DNA, pointing to generalized competitive binding or tail displacement mechanisms in nucleosome recognition by binding partners. Our study further demonstrates that post-translational modifications and mutations in histone tails can alter the tail-DNA binding modes and regulate the binding of partners to the nucleosome.

## Methods

### Construction of full nucleosome models with the native DNA sequence

There have been very few native genomic DNA sequences used in experimental and computational studies of nucleosomes. Recently we applied an integrative modeling method for constructing a high-resolution atomistic model of a yeast centromeric nucleosome with the native DNA centromeric sequence [20]. Here, we constructed a structural model of a nucleosome with the DNA sequence from a well-known oncogene, *KRAS*, which has been shown to harbor many mutations in cancer patients. In order to do this, we first identified the precise translational positioning of DNA with respect to the histone octamer. To determine the dyad position of the nucleosome, we applied a previously developed nucleosome mapping protocol to Micrococcal nuclease (MNase) experimental data using the hg19 human genome assembly [21]. Fragments of 147 bp lengths of high-coverage MNase-seq reads were used, and the dyad positions were determined as middle points of these fragments [22]. Next, we identified a well-positioned nucleosome as the first nucleosome positioned downstream of the transcription start site (the +1 nucleosome of the *KRAS* gene). To create a structural model of the full nucleosome with the DNA linkers, we used a high-resolution X-ray structure of a nucleosome core particle (NCP) formed by *Xenopus laevis* canonical core histones and human α-satellite sequence (PDB:1KX5) [23]. Then we linearly extended DNA from both ends by adding 20 bp linker segments using the NAB software (one of the H3 tails was slightly rotated to avoid steric clashes with the linker DNA by setting ψ angle of Lys36 to − 35°) [12, 24]. The native DNA sequence was selected from the human genomic region centered around the *KRAS* +1 nucleosome dyad and flanked by the 93 bp segments on each side (Figure SM1). Finally, we embedded the native DNA sequence onto the structural nucleosome model using the 3DNA program [25].

There are several structures in PDB which contain coordinates of partially resolved histone tails, which can be used in the *in-silico* studies of the nucleosomes. However, histone tails are intrinsically disordered, and their conformational ensemble covers a wide spectrum of possible configurations. Since previous studies demonstrated the rapid condensation of tails on the nucleosomal and linker DNA[11, 12], we wanted to make sure that this was not the result of initial configurations skewed toward particular conformations. Therefore, we constructed several nucleosome models with different initial tail configurations and used them for the simulations. First, we explored the existing high-resolution NCP structures (with a resolution higher than 3 Å) with the full or partial histone tail atomic coordinates in PDB [26], out of which two structures (PDB:1AOI and PDB:1EQZ) were selected based on their high resolution and partially solved histone tails. H3 and H4 tail coordinates were taken from 1AOI, and one H3 tail and two H2B tail coordinated from 1EQZ, while the conformations of other tails were taken from structure 1KX5. In those cases where templates did not contain all residue coordinates at the end of histone tails, missing residue coordinates were modeled by linearly extending existing tail conformations (dihedral angles for each residue were Φ angle = −60° and Ψ angle = 30°). As a result, two models (Model A and Model B) were built.

Furthermore, we constructed two additional models by linearly extending histone tails from the histone core into the solvent. Namely, we clipped all tails from the original 1KX5 structure at sites H3K37, H4K16, H2A A12-K118, and H2BK24 following histone tail definition from [12] and then tails were linearly reconstructed using the building structure plugin in Chimera [27] (dihedral angles used for each residues Φ = −60° and Ψ = 30°). In one initial model (Model C), tails were extended from the histone core following the backbone orientation of the last two residues at the truncated sites. We also built another initial model where histone tails were extended into the solvent symmetrically oriented with respect to the dyad axis (Model D). The Modeller software was used to remove steric clashes in tail residues surrounding the truncated sites [28]. Overall, we constructed four models with different initial tail conformations for simulations (Table SM1 and Figure SM2).

### Choice of force fields, water models, and ion parameters

An appropriate choice of the force field, water model, and ion parameters is required to simulate highly charged large macromolecular systems such as nucleosome, to model protein-DNA interactions, conformations of disordered histone tails, and nucleosome interactions with the solvent and ions. In order to achieve sufficient and accurate sampling of histone tail conformations, we explored and compared different force fields, water models, and ion parameters in our simulation protocols.

CHARMM and AMBER force fields are both widely used in the simulations of macromolecules and are being continuously improved [29–31]. Here, we selected two sets of recently developed protein and DNA force fields in CHARMM and AMBER packages. One is CHARMM36m for protein and CHARMM36 for DNA force fields, where recent developments in CHARMM36m have shown improvements in generating the conformations for intrinsically disordered proteins (IDPs) [30]. Another set includes the AMBER ff14SB force field for protein and OL15 force field for DNA [29–32]. Regarding water models, TIP3P is a 3-site rigid water model, widely used in many studies. Although the TIP3P water model was shown to offer a relatively good compromise between speed and accuracy, it poorly reproduced the physical properties of water and, in some cases, had large disagreements with experiments in terms of binding free energy calculations in protein-ligand systems [33, 34] and modeling of conformational ensembles of IDPs [35]. An Optimal Point Charge (OPC) water model is a 4-point rigid water model, which reproduces comprehensive sets of water liquid bulk properties and delivers noticeable accuracy improvement in simulations of RNA, thermodynamics of ligand binding, small molecule hydration, and intrinsically disordered proteins [36–38]. Most recently, the OPC water model, together with the AMBER force field, offered remarkable improvements over the TIP3P water model in the modeling of the conformational ensembles of IDPs [39]. Here, we applied the TIP3P water model with the CHARMM force field and AMBER force field with the OPC water model. Regarding the choice of ion parameters, we selected the 12-6 HFE parameter set for monovalent ions, developed for the OPC water model [40], and Beglov and Roux ion parameters [41] were used with the CHARMM force field and TIP3P water model.

For four constructed nucleosome models (Model A, B, C and, D), we performed simulations using the AMBER and CHARMM force fields. For the AMBER simulations with the OPC water model, for each nucleosome model, we performed five independent runs with different seeds, four runs had 200 ns simulation time, and one run reached 2500 ns for the purpose of observing phenomena on a longer time scale. For model D (nucleosome model with the symmetrically extended tails), we performed two 2500 ns simulation runs using GROMACS with the OPC water model and AMBER force field. In parallel, we performed three 100 ns simulations for each nucleosomal model using the CHARMM force field and the TIP3P water model. A summary of all simulation runs for histone tail sampling is provided in Table SM 1.

### Simulation protocols

The MD simulations using the AMBER force field and OPC water model were prepared and performed with the Amber18 package [42] and GROMACS version 2019.3 [43]. MD simulations using the Amber18 package (20 simulations runs in total) were performed as following (Table SM1). Nucleosome structures were solvated with 0.15 M NaCl in a cubic water box with at least 20 Å from the protein to the edge of the water box (detailed information is provided in Table SM1). Systems were maintained at T=310 K using the Langevin dynamics with the integration step of 2 fs and collision frequency γ = 1 ps^−1^. The Berendsen barostat was used for constant pressure simulation at 1 atm. SHAKE bond length constraints were applied for bonds involving hydrogens. The cut-off distance for non-bonded interaction calculations was 10 Å. Particle Mesh Ewald (PME) method with a spacing of 1 Å and real space cut-off 12 Å was applied for the electrostatic calculations. Periodic boundary conditions were used, and the trajectories were saved every 20 ps. All systems were first subjected to 10,000 steepest descent minimization and then for another 10,000 conjugate gradient minimizations. After minimization, systems were gradually heated from 100 K to 310K in the NVT ensemble and then switched to the NPT ensemble for 500 ps equilibrations before production runs.

Two simulation runs using the GROMACS package were performed as following (Table SM1). A cut-off of 10 Å was applied to short-range non-bonded interactions, and the Particle Mesh Ewald (PME) method was used in calculating long-range electrostatic interactions. Long-range dispersion corrections for energy and pressure were applied for long-range Van der Waals interactions. Covalent bonds involving hydrogens were constrained to their equilibrium lengths using the LINCS algorithm. The solvated systems were first energy minimized using steepest descent minimization for 10,000 steps, gradually heated to 310 K over the course of 800 ps using restraints, and then equilibrated for a period of 1 ns. After that, the production runs were carried out in the NPT ensemble up to 2.5 μs, with the temperature maintained at 310 K using the modified Berendsen thermostat (velocity-rescaling) and the pressure maintained at 1 atm using the Parrinello–Rahman barostat.

Simulations using CHARMM forcefield and TIP3P water models were prepared with VMD [44] and performed using NAMD 2.12 package [45]. The simulation protocol was similar to the previous one. The systems were initially subjected to 1000 steps of energy minimization with all protein and DNA atoms fixed and then to another 10,000 steps of minimization without constraints. Next, we performed four rounds of 200 ps equilibrations with elastic constraints on C-α atoms of protein and P atoms of DNA backbone, which were gradually relaxed as follows: 90 -> 45-> 9-> 0 kcal/mol/A^2^.

### Simulations of nucleosomes with mutated and post-translationally modified histones

To elucidate the effects of mutations and histone modifications on tail-DNA interactions, we performed multiple sets of simulations, including lysine acetylation, lysine methylation, serine/threonine phosphorylation, and Arg -> Ala substitutions introduced at the same time or at one residue at a time (Table SM 2). AMBER force field and OPC water model were applied using protocols described above. Mutations and PTMs were introduced to nucleosome structures with LEaP in the AMBER package [42], and locations and force field parameters of PTMs were taken from previous studies [46, 47]. For each set of simulations, we performed five independent runs with different random seeds, of which four runs had 200 ns simulation time and one run of 1,000 ns.

### Trajectory analysis

We only used trajectories from simulations with the OPC water model for analyses. Trajectories were visualized and analyzed using a set of TCL and Python scripts that utilized the capabilities of VMD [44], 3DNA [25], and AMBER Tools [42]. The trajectory frames were superimposed onto the initial models by minimizing RMSD values of *C*^*α*^ atoms in histone cores (Table SM3). In the analysis of histone tail-DNA interactions, tail-DNA atomic contacts were calculated for trajectory frames of every 1 ns. The first 200 ns frames of each 2000 ns run and 50 ns frames of each 200 ns run were disregarded as an initial conformational equilibration period. The contacts of atoms between histone and DNA were defined between two non-hydrogen atoms located within 4 Å. For each DNA base pair, we calculated the mean number of bound histone tail heavy atoms averaged over frames. Then, we defined the histone tail preferred binding regions as DNA base pairs that had more than five contacts on average with histone tails.

The residence time of the histone tail was defined as the time during which tails remained bound to DNA in the simulations. Two types of residence time were calculated: individual residue residence time (*τ*_*r*_) and the full tail residence time (*τ*_*f*_). Here, a bound state for an individual residue was defined if at least one heavy atom of a residue had a contact with DNA. An unbound state for the full tail was defined if less than a certain fraction of histone residues maintained contacts with the DNA molecule (different values of this threshold were tested; see Supplementary Materials). Since full histone tails undergo very rapid fluctuations before retaining stable binding with DNA during the simulations, we ignore *τ*_*f*_ of shorter than 10 ns. DNA solvent accessible surface area (SASA) was calculated using VMD [44] with a probe distance of 1.4 Å for every 1 ns frames. The nucleosomal and linker DNA solvent accessibility area change upon histone tail binding was calculated as the difference between the SASA of DNA with tails bound to it and without tails.

The binding free energy between histone tails and DNA was calculated using the MM/GBSA (Molecular Mechanics Generalized Born Surface Area) method implemented in the Amber18 package. We performed calculations for every 1 ns frame (ignoring the first 50 ns in the 200 ns trajectories and 200 ns in 2000 ns trajectories), and residue-wise decomposition was applied to derive the binding energy per tail residue. Each copy of a tail within a simulation was considered as a separate observation of the tail ensemble. Thereby there were two conformations per frame per histone type. In all calculations, the standard error (SE) of the mean from independent simulation runs (22 runs in total) were estimated.

### Analysis of experimental structures of nucleosomal complexes

We extracted all nucleosome complex structures from PDB [48] for our analysis of nucleosome-binding proteins and then removed 20 structures that did not contain the complete histone octamer or had extensive DNA unwrapping or sliding along the octamer (structures where proteins interacted with the linker DNA, were kept in our analysis). The interaction between a nucleosome and a binding partner was defined if histone proteins and/or nucleosomal and/or linker DNA had at least one non-hydrogen atom within 4 Å of nucleosome binding proteins. Functional classifications of nucleosome binding proteins were performed using the general protein function annotations from UniProt [49]. To quantitively characterize the degree of DNA interfacial overlap between DNA-histone tails and DNA-nucleosome binding proteins, we calculated the fraction of interface overlap as a number of DNA base pairs found on both DNA-tail (from MD simulations) and DNA-partner binding interfaces (from PDB experimental structures) divided by the number of DNA base pairs making contacts with nucleosome binding proteins in a PDB structure.

## Results

### DNA binding properties differ between histone tail types

Histone tails have high conformational flexibility, and their conformational sampling represents a major challenge. To address this problem, we have built four nucleosome models with different initial histone tail configurations and performed 34 different runs totaling in about 20 microseconds of simulations (Table SM1), which can provide a quite extensive overview of the histone tails’ conformational and interaction landscape. In concordance with other *in silico* and experimental studies [11, 12, 14, 50, 51], we observe a relatively rapid condensation and extensive interactions of histone tails with the nucleosomal and linker DNA. The dynamics and kinetics of histone tail binding depend on the choice of water model and, to a lesser extent, on a choice of the force field. Using the OPC water model, we observe a slower histone tail condensation on the nucleosomal and linker DNA on the timescale of ~100 ns compared to 10-50 ns using the TIP3P water model. One possible reason for these differences could arise from self-diffusion coefficient values used for the TIP3P water model being about 2.5 times larger than the experimental values, which could artificially accelerate the motions in the simulations. Simulations using the OPC water model show many rapid interconversions between tail-DNA bound and unbound states, pointing to a more dynamic histone tail behavior compared to simulations with the TIP3P water model where histone tails remain in the bound state with DNA most of the time [11, 19] (Figure 1a). Since the TIP3P water model can over-stabilize the compact states compared to extended conformations [39], histone tails are rarely observed in unbound states after initial condensation [11, 12]. Therefore, thereafter we only use simulations with the OPC water model for analyses.

**Figure 1.**
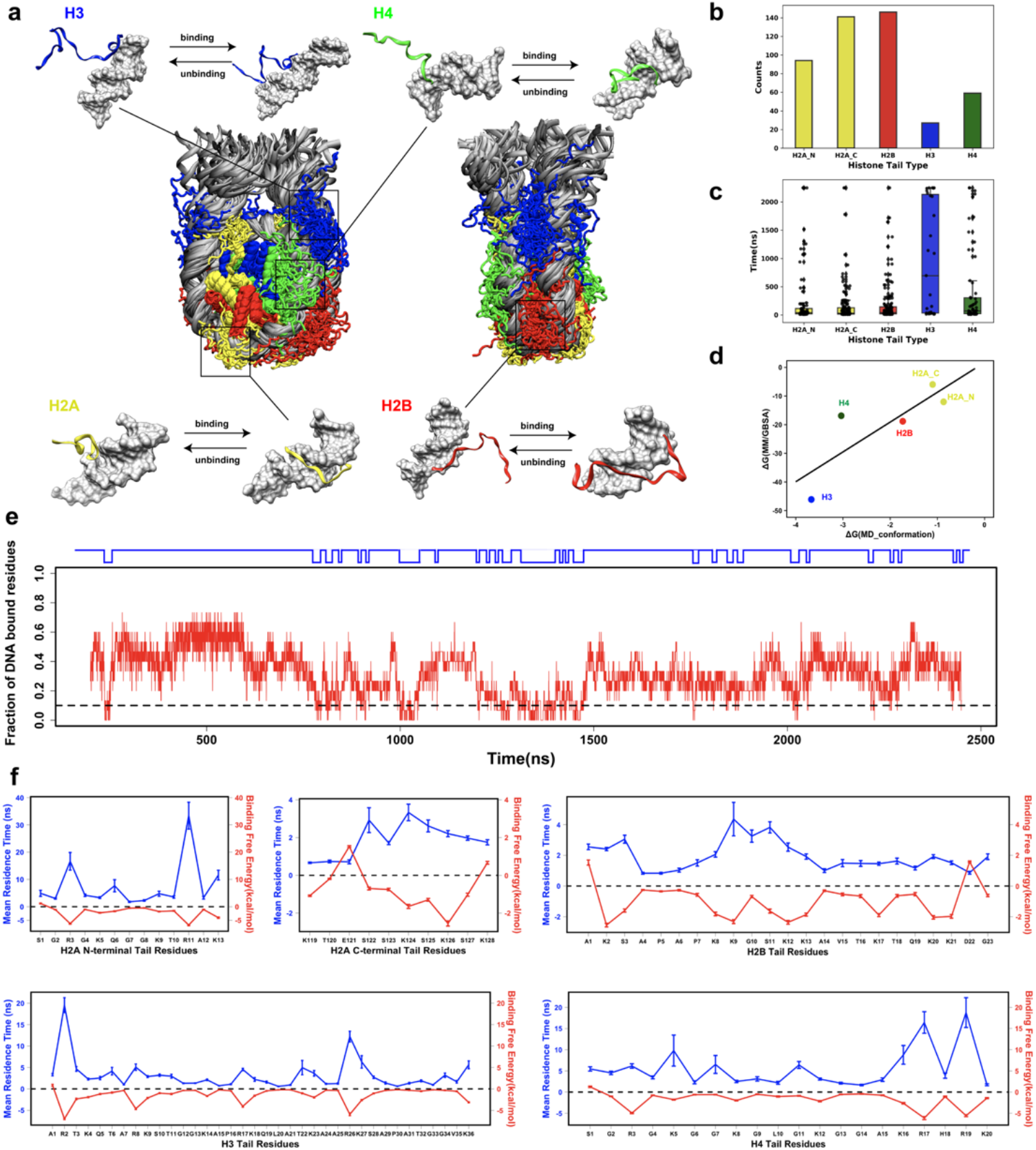
Binding of histone tails to nucleosomal and linker DNA in the context of the full nucleosomes. a) Interconversions of DNA-bound and unbound tail conformations. The conformational snapshots are taken from the last frame of each simulation run and superimposed onto the initial models by minimizing RMSD values of *C*^*α*^ atoms in histone cores. b) A total number of full histone tail binding/unbinding events observed in all simulations for both copies of histones. c) Full histone tail residence time. Each point represents a binding/unbinding event observed in simulations for either histone copies. Residence time shorter than 10 ns is excluded as this time is required for establishing stable interactions with DNA. An unbound state for the full tail is defined if less than 10% of the tail residues maintain contacts with the DNA (different values of this cut-off have been tested, see Supplementary Materials). d) The correlation between the histone tail-DNA binding free energy (ΔG) estimated by two independent approaches: derived from counting the number of bound/unbound frames in histone tail conformational ensemble (number of binding/unbinding events >=5, Table SM4) and estimated by MM/GBSA approach (Table SM5). e) A representative run shows a fraction of DNA bound residues and tail binding/unbinding events during simulation for H2B tails (see Figures SM13, 14, 15, 16, 17 for other tails). f) Individual residue residence time (*τ*_*r*_) and binding free energy estimated by MM/GBSA approach. Residence time and binding energies are averaged over 1 ns frames, and the error bars represent standard errors of the mean calculated from independent simulation runs. We ignore *τ*_*r*_ of H2AK13 calculated from one simulation run of Model B, where H2AK13 does not unbind from DNA during the simulation.

To further characterize the kinetics of histone tail-DNA interactions, we count a total number of transitions from unbound to bound states and compute histone tail residence time to estimate the effective time that histone tails stay bound to the DNA molecule (as the inverse of the rate constant, *τ* = 1/*k*_*off*_), evaluating full tail residence time (*τ*_*f*_) and individual residue residence time (*τ*_*r*_). As can be seen in Figure 1b and c, the number of binding-unbinding events and residence time varies considerably between histone types. H3 has the highest residence time among all tails, up to two microseconds, and is characterized by relatively fewer unbinding events. It is followed by H4 and H2A N-terminal tails, whereas H2A C-terminal and H2B tails have the lowest average residence time and lower binding free energy per tail residue with DNA compared to other tails (Figure 1c, Table SM 4 and 5). ANOVA analysis and Tukey HSD test confirm that H3 and H4 tails have significantly different residence times compared to the residence time of other tails (Table SM6). To characterize the binding kinetics in more detail, we calculate the individual residue residence time (*τ*_*r*_) (Figure 1e), which is found to be on the time scale of several to tens of nanoseconds, demonstrating very rapid and frequent transitions between bound and unbound states and jittery conformational rearrangements of histone tails in the bound state. Congruent with these findings, residues with long *τ*_*r*_ have a high binding free energy with DNA (Figure 1e). We further compare our estimates of dissociation constants from counting the number of binding/unbinding events with the binding free energy estimates coming from a set of independent MM/PBSA calculations (Table SM 4,5 and Figure SM7). Overall, it shows a remarkable linear association with R^2^ = 0.9, pointing to a reasonable conformational sampling of histone tails’ binding performed in our study (Figure 1d).

As was observed in many previous studies, one of the most prevalent modes of interaction between histone tails and DNA was the insertion of the arginine and, in some cases, lysine side chains into the DNA minor and major grooves serving as anchors stabilizing these interactions [12, 52]. Figure 1e shows that anchoring of arginine is indeed critical in determining the tail’s longest residence time. H2A C-terminal and H2B tails do not have arginine residues and therefore exhibit the shortest *τ*_*f*_, while H3 and H4 tails are arginine-rich and have the longest *τ*_*f*_. For tails without arginine residues, the most prominent mode of interaction is between lysine and serine residues and DNA.

### Histone tail dynamics modulate the nucleosomal and linker DNA accessibility

Interactions between histone tails and DNA may decrease their respective solvent accessibility. At the same time, upon unbinding, histone tails and DNA become more accessible to nucleosome-binding proteins [11]. We analyze the interaction modes of histone tails and estimate the changes of nucleosomal and linker DNA solvent accessibility imposed by the tail binding. Due to the 2-fold pseudo-symmetry of the nucleosome structure, upon exhaustive conformational sampling, one should expect that each histone copy samples a similar phase space region (Figure SM8). Indeed, we show that there is a significant correlation coefficient between the mean number of tail-DNA contacts occupied by both copies of histone tails (Figure SM9). Therefore, below we report a combined conformational ensemble from both copies of histone tails. As can be seen in Figure 2a, tails of different histone types preferably interact with the specific DNA regions. H2A N-terminal tails bind to the nucleosomal DNA at SHL ±4, whereas H2A C-terminal tails are mostly bound at SHL ±7 and near the dyad. Interaction modes of H2B tails encompass a more extensive DNA binding interface compared to other tails. The relatively short residence time of H2B tails identified in the previous section suggests a dynamic behavior of H2B tails allowing them to search a large surface area on DNA without being kinetically trapped in specific conformations. Being the longest, H3 tails can also interact with DNA in multiple regions with the longest residence time: near the dyad, at SHL ±6 to ±7 as well as with the linker DNA. Finally, the preferred interaction mode of H4 tails is near SHL ±1 and ±2.

**Figure 2.**
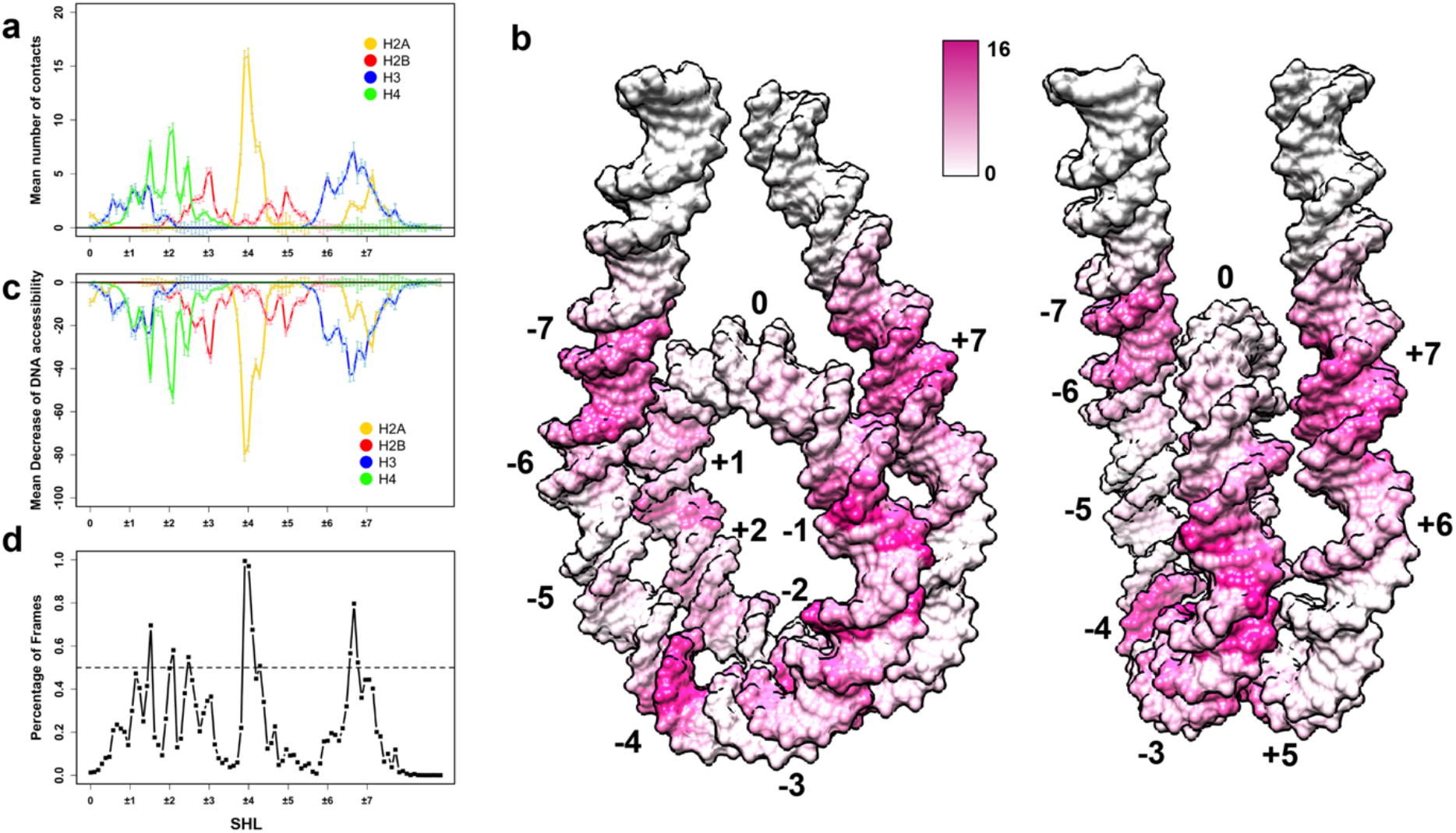
Nucleosomal and linker DNA solvent accessibility, SASA, modulated by histone tail binding. a) Mean number of contacts between histone tails and nucleosomal/linker DNA averaged over all independent simulation runs plotted in the DNA coordinate frame, zero corresponds to the dyad position and SHL locations are shown as integers; a combined conformational ensemble from both copies of histone tails is shown. b) Mean number of contacts between histone tails and DNA mapped onto the molecular surface of the nucleosomal and linker DNA. c) Changes of DNA solvent accessibility imposed by tail binding averaged over 1ns frames in Å^2^ units. d) Percentage of frames with more than 25% SASA decrease upon tail binding. The percentage of accessibility change for a DNA base pair is defined as a difference between SASA of nucleosomal/linker DNA with and without bound tails divided by total SASA. The error bars represent standard errors of the mean calculated from independent simulation runs.

In summary, we identify those DNA regions which are in tail bound states most of the time (more than in 50% MD frames) and have more than 25% solvent accessible surface area decrease - those regions include SHL±1.5, ±2, ±2.5, ±4, and ±7 (Figure 2) and the solvent accessibility of these regions is directly affected by the tail binding (Figure 2c, Figure SM 10,11). The change of the DNA solvent accessible surface area is correlated with the number of contacts between DNA and tails. Some DNA regions that interact with histone tails undergo a substantial decrease of SASA, whereas the average decrease in SASA per one base pair is up to 80 Å^2^(Figure 2c).

### Histone tails and nucleosome-binding proteins target overlapping regions on nucleosomal/linker DNA

Nucleosomes, being the hubs in epigenetic signaling pathways, are targeted by a wide spectrum of nucleosome-binding proteins that interact with the specific regions on nucleosomal/linker DNA and histones [53]. To this end, we perform a systematic analysis of interaction modes of nucleosome-binding proteins using available nucleosome complex structures in PDB [48], totaling in 89 structures (Figure 3a). The functional classification of nucleosome-binding proteins shows that the majority of them include chromatin remodelers and transcription regulatory proteins. More than 70% of nucleosome-binding proteins recognize some part of DNA molecules, and most of them exhibit multivalent binding modes interacting with both histones and DNA. Among multivalent interactors, about half of them recognize histone tails as well as DNA (H3 and H4 tails, Table SM7), and another half recognizes DNA and histone core residues. An example of chromatin remodeler ISWI, which binds to nucleosomal DNA at SHL ±1.5 and H4 tails, is shown in Figure 3f.

**Figure 3.**
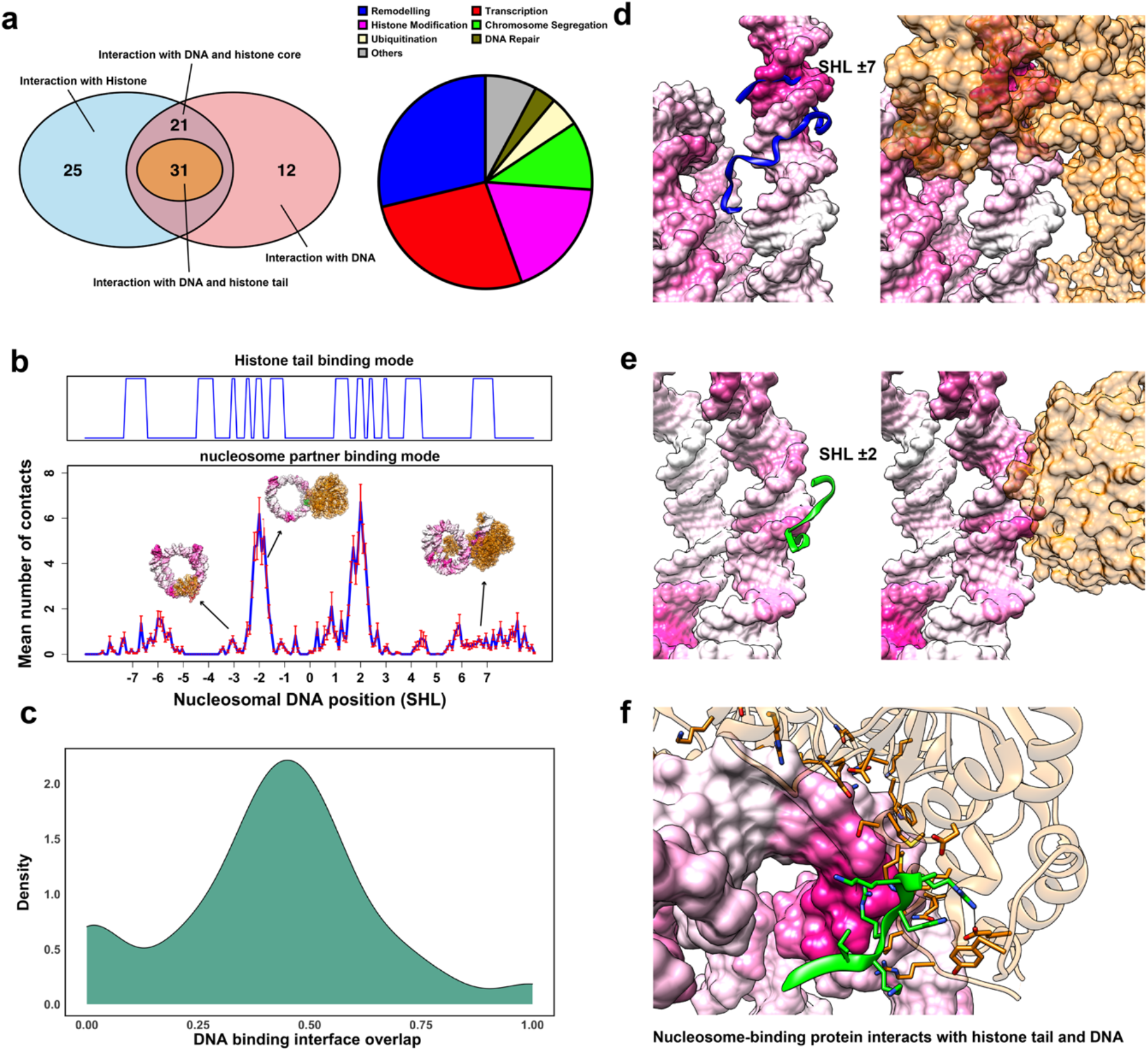
Histone tail-modulated recognition modes of nucleosomes by binding proteins. a) A summary of available nucleosome complex structures classified based on their binding entity and function. b) Mean number of contacts between the nucleosomal/linker DNA and nucleosome-binding partner (averaged over all complexes) plotted in DNA coordinate frame. The error bars represent standard errors calculated from the number of contacts of different nucleosome complex structures. Top track shows the presence or absence of five or more contacts between histone tails and DNA regions. c) An overlap between tail-DNA and partner-DNA binding interfaces. It is calculated for each nucleosome complex structure and the distribution is smoothed using kernel density estimation. d), e) Examples of INO80 chromatin remodeler (PDB: 6HTS) and UV-damaged DNA-binding protein (PDB: 6R8Z) targeting overlapping regions on DNA. Histone tail representative conformations are taken from simulations and superimposed onto the PDB structures. The intensity of the color of DNA surface is scaled with the mean number of contacts between histone tails and DNA as in Figure 2b. f) An example of chromatin remodeler ISWI which binds to both histone H4 tail and DNA (PDB: 6IRO). Here coordinates of histone tails are taken from the PDB structure. Nucleosome-binding proteins are colored as orange and histone tails are colored using their canonical colors.

As can be seen in Figure 3b, nucleosome-binding proteins show distinctive preferred binding regions on DNA around SHL ±1, ±2, SHL± 6 and SHL±7 and to a lesser extent on linker DNA and near the nucleosome dyad. If we compare these interfaces to the preferred interaction modes of histone tails on DNA observed from our simulations (see previous section), it is clear that there is a considerable interface overlap at SHL ±1, ±2, and ±7 (Figure 3b). Namely, dynamic histone tails and many nucleosome-binding proteins seem to target overlapping and mutually exclusive regions on nucleosomal or linker DNA. For each nucleosome complex structure where a binding partner interacts with DNA (64 complexes), we calculate a fraction of DNA interface shared between the histone tail ensemble (from MD simulations) and the nucleosome-binding proteins (from structures) (Figure 3c) and find that in 89% of them (57 complexes) interfaces are mutually exclusive (at least one base pair shared on binding interfaces). Figures 3d,e show the chromatin remodeler, INO80, bound at SHL ±7 and to the linker DNA, and the UV-damaged DNA-binding protein bound at SHL ±2. These DNA regions can also be occupied by H3 and H4 histone tails, as evident from the tail ensemble of MD simulations.

### Histone tail post-translational modifications alter tail-DNA interactions

Next, we try to elucidate the roles of PTMs and mutations in modulating the histone tail-DNA binding modes. Histone tails harbor different PTMs that can affect histone tail dynamics and interactions in the context of the full nucleosome. In addition, histone genes are mutated in many cancers and might represent oncogenic drivers [54]. We perform alignments of all histone protein sequences and then map nucleosome binding sites (using all collected nucleosome complex structures from PDB) and histone cancer missense mutations from a recent histone mutation dataset onto them [54–56]. As can be seen in Figure 4a, many of cancer mutations affect the charged residues in histone tails and alter the tail post-translational modification sites. To further elucidate the effects of PTMs and mutations on tail-DNA interactions, we systematically compare tail-DNA interaction modes for unmodified tails and for various types of modified tails (lysine acetylation, lysine tri-methylation, serine/threonine phosphorylation, and Arg->Ala mutations) by performing simulations in the context of the full nucleosome (Table SM2). Here we estimate the maximal possible effects of such modifications as these sites might not be modified at the same time in a cell.

**Figure 4.**
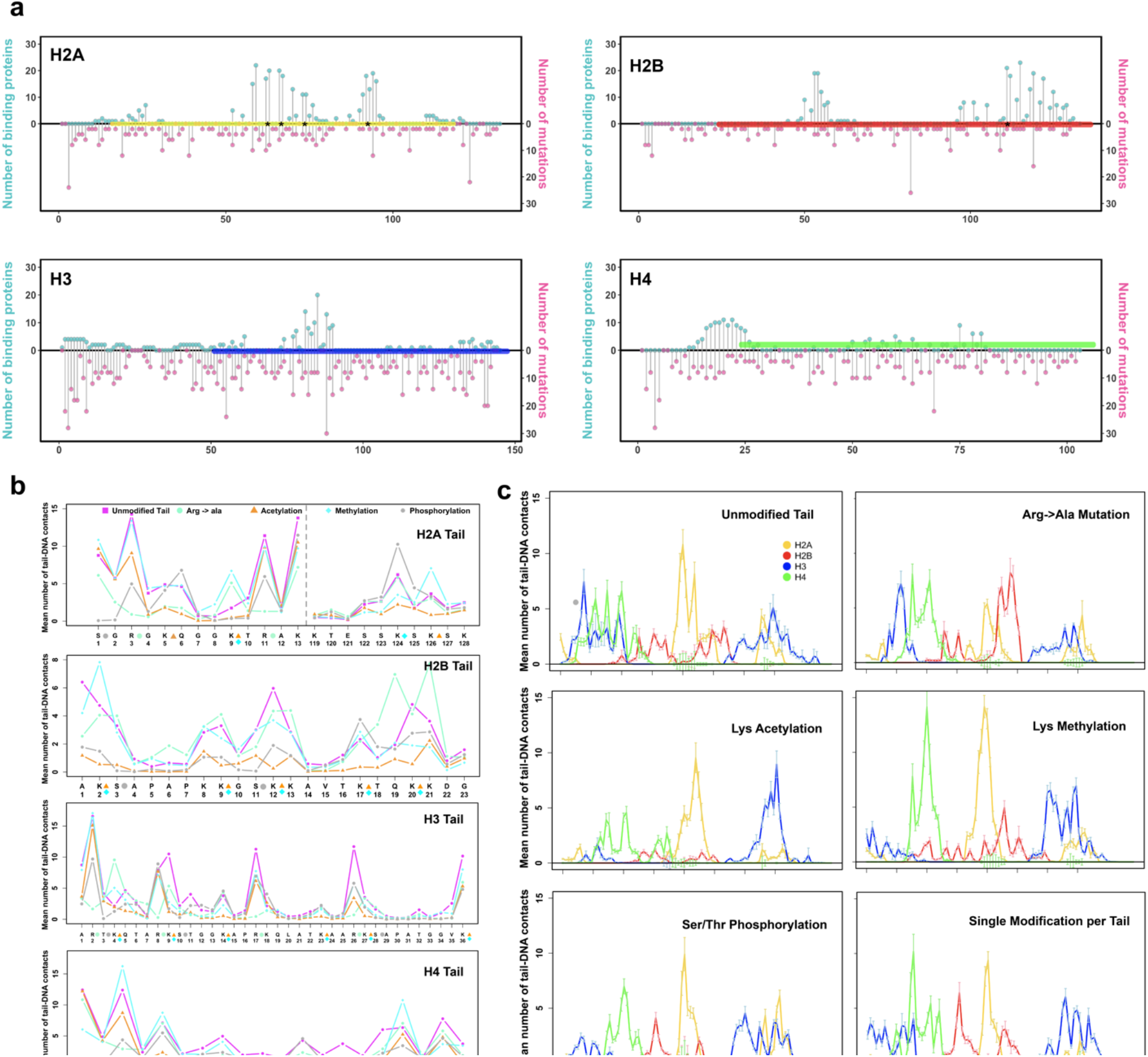
Histone tail post-translational modifications and cancer-associated mutations modulate histone tail-DNA interaction modes and DNA accessibility. a) Numbers of cancer-associated mutations (pink) and binding proteins (light blue) mapped on tracks representing consensus sequence of the full alignment of histone sequences (see Figures SM18, 19, 20, 21). Globular domains are indicated as yellow, red, blue and green bars per histone type. Black asterisks denote the acidic patch residues. b) Mean number of histone tail-DNA contacts for different types of modifications. Simulations of *model D* (Nucleosome with symmetrically extended histone tail configurations) with and without corresponding modifications are shown in different colors and modifications are shown by a symbol next to the residue. c) Mean number of nucleosomal and linker DNA contacts with histone tails. For each type of modification, the reported values are averaged over 1 ns frames and the error bars represent the standard errors of the mean calculated from independent simulation runs. The locations of modifications and mutations are listed in Table SM2).

There are two main striking observations evident from Figure 4b,c. First, modifications changing the effective positive charge of the residue (lysine acetylation, serine phosphorylation, and Arg->Ala mutations) significantly affect the interactions of tails with DNA (Figure 4b), overall decrease the tail-DNA binding free energy and decrease full tail residence time *τ*_*f*_ (Table SM8, SM22). However, the amplitude of these effects depends on histone type, position of PTM in a sequence, and modification types of the residue and other surrounding residues. The most dramatic effects are observed for modifications of H4 tail: even a single charge-changing modification considerably decreases its interactions with DNA, especially near the dyad and SHL ±2 regions (Figure 4c). The effects on H3 tail dynamic behavior are more complex: although an overall number of contacts with DNA does not change much, modifications induce the redistribution of contacts: tri-methylation of H3 and acetylation of H3 introduced once at a time lead to the loss of the contacts with DNA near the dyad region and increase in the number of contacts with the DNA near the exit site, SHL ±6,7. Second, our findings point to the crosstalk between different modified sites so that a modification in one site may lead to substantial changes of interactions with DNA in another histone site. For example, the number of contacts of H3K4 with DNA almost triples when the interactions of H3R2 are suppressed through an Arg ->Ala mutation. Tri-methylation of H4K5 enhances the interactions of H4R3 with DNA, whereas the interactions of H3R26 with DNA are suppressed by phosphorylation of H3S28.

## Discussion

Nucleosomes are elementary building blocks of chromatin and, at the same time, may act as signaling hubs by integrating different chromatin related pathways [53] and directly participating in the regulation of many epigenetic processes pertaining to the access of chromatin factors to DNA and histones [53]. It has been long debated about how the DNA solvent accessibility and mutability can be modulated for those regions which are packed in nucleosomes [6]. According to the commonly used static model, the DNA accessibility follows the ten base pair periodicity patterns of rotational positioning of nucleosomal DNA [20]. However, we have shown that there is another important layer in this mechanism, which stems from the histone tail dynamics. Even though histone tails extensively condense on the DNA, comprehensive simulations performed in this study allow us to observe many histone tail binding and unbinding events. Namely, we demonstrate that the tails undergo rapid transitions between bound and unbound states, and the kinetics of these processes depend on the histone type. The interactions between tails and DNA are transient, and switching between tail conformations occurs on the time scale from tens to hundreds of nanoseconds in the form of jittery motions, with H2A C-terminal, H2A N-terminal, and the H2B tails having the shortest residence time on DNA and H3 and H4 tails having the longest residence time.

The interactions of histone tails with the DNA molecule within the same nucleosome affect the nucleosomal and linker DNA accessibility - even though the interactions of individual tails with DNA are transient, DNA regions SHL±1.5, ±2, ±2,5, ±4, and ±7 are occluded from the solvent by different types of histone tails most of the time. Histone tail interactions with the DNA may modulate the accessibility of both the DNA and histone tails themselves to other binding biomolecules. Indeed, it has been shown for PHD1/2 readers that they can bind up to ten-fold tighter to histone peptides compared to free nucleosomes [19]. A recent large-scale experimental study also demonstrates that many chromatin factors show enhanced binding to tailless nucleosomes compared to the full nucleosome, resulting from the increased solvent accessibility of DNA [57]. Our estimates of the binding free energy of the histone tail binding to DNA based on conformational sampling are on the order of several kcal/mol for H3 and H4 tails with the strongest binding exhibiting for H3 tail (Table SM4), which is consistent with previous experimental data on the tail-DNA binding free energies [58]. Similar effects, although not investigated here, may pertain for long tails spanning the distance between the neighboring nucleosomes.

Our second prediction, based on the analysis of dynamic MD ensemble of histone tail conformations and nucleosome experimental structural complexes, indicates that nucleosome-binding proteins and histone tails may target overlapping and mutually exclusive regions on nucleosomal or linker DNA around SHL ±1, ±2, and ±7. This trend is observed for 89% of studied complexes and may point to a possible competitive binding mechanism: nucleosome-binding proteins compete with DNA if they recognize tails and compete with histone tails for binding to DNA (Figure 5). The competition between chromatin factors has been previously recognized as a major determinant of various chromatin states [59]. At the same time, our analysis identifies 31 nucleosome-binding proteins interacting with both histone tails (via H3 and H4 tails) and nucleosomal or linker DNA (Table SM7 and Figure SM12) and importantly, in those complexes, histone tails do not have direct contacts with DNA. Such recognition patterns could be explained by the multivalent binding and/or by a recently proposed tail displacement model (Figure 5) [60, 61]. According to the tail displacement model, interactions of DNA-binding domains (DBD) of a nucleosome-binding protein with the nucleosome can displace histone tails from their DNA preferred binding modes. It makes them more accessible for recognition by reader domains (Figure 5). This is supported by recent studies showing that H3 tail-DNA interactions inhibit the activity of histone-modifying enzymes [15] and binding of BPTF PHD finger to tails [11]. The displacement of histone tails, in turn, can be facilitated by the competitive binding between histone tails and DBDs if they both recognize the same regions on DNA. This could accelerate the unbinding of tails from DNA and enhance the recognition of tails by the reader domains. Such competitive binding or tail displacement processes are controlled by local concentrations of tails and binding partners and by their binding affinities, the latter was experimentally determined to be ~8-10 kcal/mol, on the same order of magnitude as the strength of tail-DNA interactions estimated in our study (Table SM9) [62].

**Figure 5.**
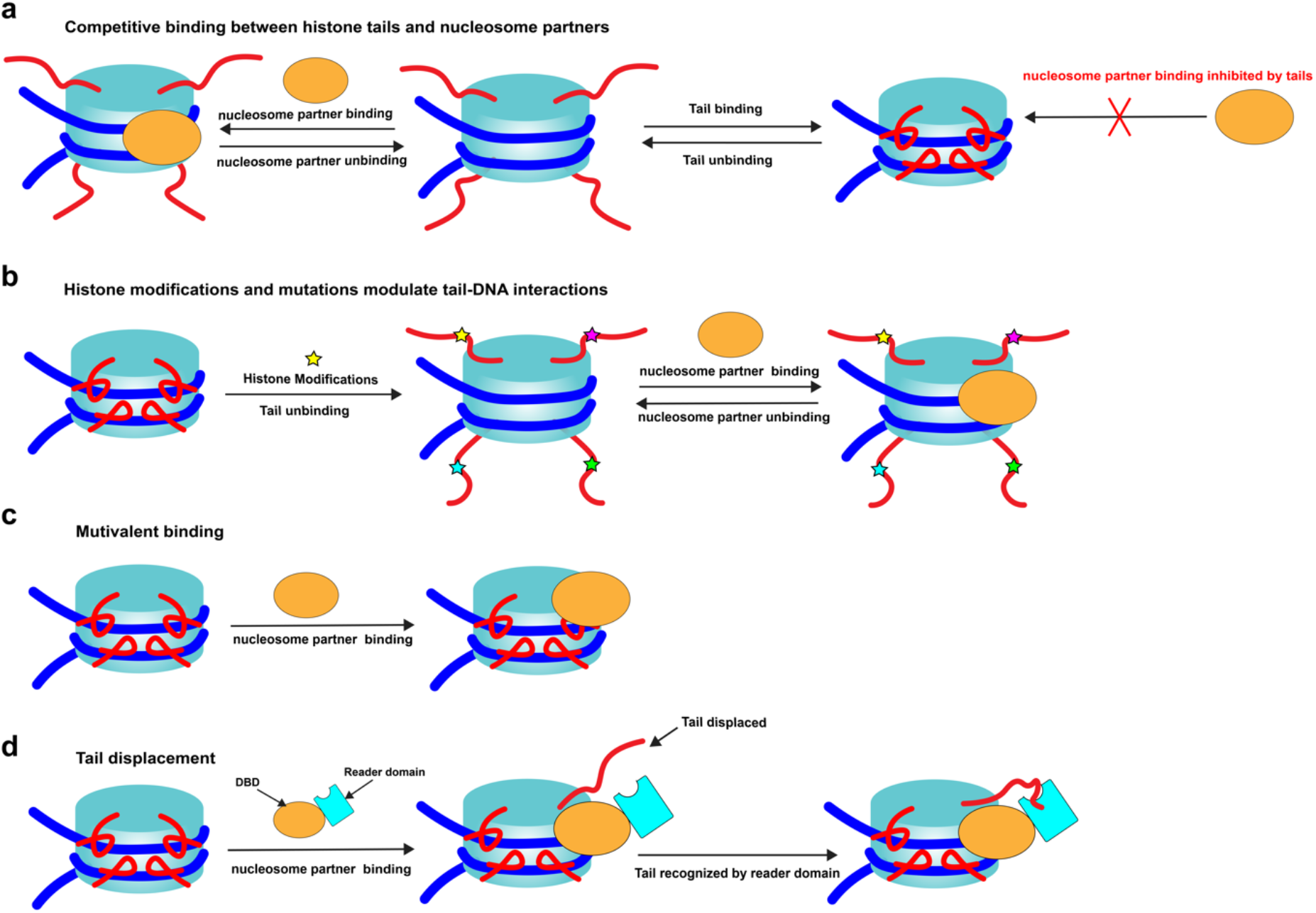
A generalized model explaining how tails, their mutations, and post-translational modifications can modulate nucleosomes’ interactions with nucleosome-binding proteins. a) Histone tails’ interactions with the DNA modulate the accessibility of the DNA to other binding proteins; Nucleosome-binding proteins compete with histone tails for binding to DNA. b) Charge-altering modifications and mutations in histone tail residues can suppress tail-DNA interactions, enhance histone tail dynamics, and regulate binding of proteins to nucleosome. c) Nucleosome-binding proteins exhibit multivalent binding modes and recognize both histones and DNA. d) Reader domains of nucleosome-binding proteins compete with DNA if they recognize tails; Interactions of DNA-binding domains (DBD) with nucleosome can displace histone tails from their DNA preferred binding modes and increase their accessibility for recognition by reader domains.

Histone tail post-translational modifications can be responsible for the regulation of tail-DNA interactions through the alteration of histone tail binding modes (Figure 5). As we demonstrate, neutralizing charge-altering modifications and mutations in histone tail residues overall may suppress tail-DNA interactions and enhance histone tail dynamics. Consequently, this mechanism can boost the interactions between nucleosomes and nucleosome-binding proteins, which specifically recognize certain histone tail sites. Consistent with these observations, phosphorylation and acetylation of H3 tails were found in recent studies to weaken H3 tail-linker DNA interactions to stimulate the H3 tail dynamics [11, 15]. We show that histone modifications may have local or long-distance effects, and modification in one site can influence the dynamics and histone-DNA interactions in another site. As an example, interactions of arginine residues with DNA can be modulated by tri-methylation of lysine located up to several residues apart in sequence.

We argue here that histone tails are crucial elements in coordinating the transient binding and recognition of different chromatin factors to nucleosomes and thereby contribute to the regulation of epigenetic processes in time and space. Their disordered dynamic nature is a prerequisite for this task allowing histone tails to bind to different partners via the same interface with high or low affinity and high specificity. Similar to well-documented cases of the disorder-mediated control of the exposure of protein-protein interfaces, here we argue that akin mechanism can pertain to protein-DNA interface exposure at the level of the nucleosome and show that modulating DNA access through histone tails might represent a rather general mechanism. The quantitative characterization of these dynamic processes has been very challenging, and data is still largely lacking. The future focus on the development of experimental and computational techniques elucidating the spatial and temporal hierarchy of dynamic chromatin processes may close this gap in our understanding.

## Acknowledgements

YP and DL were supported by the Intramural Research Program of the National Library of Medicine at the U.S. National Institutes of Health. ARP was, in part, supported by the Intramural Research Program of the National Library of Medicine at the U.S. National Institutes of Health. SL and ARP were supported by the Department of Pathology and Molecular Medicine, Queen’s University, Canada. AO was supported by the funding from the National Science Foundation, MCB-1715207. ARP is the recipient of a Senior Canada Research Chair in Computational Biology and Biophysics and a Senior Investigator Award from the Ontario Institute of Cancer Research, Canada. This study utilized the high-performance computational resources from the Biowulf cluster at the National Institutes of Health (https://hpc.nih.gov/systems/) and Compute Canada (https://docs.computecanada.ca).

## References

1. Luger, K., et al., Crystal structure of the nucleosome core particle at 2.8 A resolution. Nature, 1997. 389(6648): p. 251–60.

2. Ferreira, H., et al., Histone tails and the H3 alphaN helix regulate nucleosome mobility and stability. Mol Cell Biol, 2007. 27(11): p. 4037–48.

3. Iwasaki, W., et al., Contribution of histone N-terminal tails to the structure and stability of nucleosomes. FEBS Open Bio, 2013. 3: p. 363–9.

4. Valieva, M.E., et al., Stabilization of Nucleosomes by Histone Tails and by FACT Revealed by spFRET Microscopy. Cancers (Basel), 2017. 9(1).

5. Korolev, N., A.P. Lyubartsev, and L. Nordenskiold, A systematic analysis of nucleosome core particle and nucleosome-nucleosome stacking structure. Sci Rep, 2018. 8(1): p. 1543.

6. Klemm, S.L., Z. Shipony, and W.J. Greenleaf, Chromatin accessibility and the regulatory epigenome. Nat Rev Genet, 2019. 20(4): p. 207–220.

7. Allahverdi, A., et al., The effects of histone H4 tail acetylations on cation-induced chromatin folding and self-association. Nucleic Acids Res, 2011. 39(5): p. 1680–91.

8. Bohm, L. and C. Crane-Robinson, Proteases as structural probes for chromatin: the domain structure of histones. Biosci Rep, 1984. 4(5): p. 365–86.

9. Zhou, B.R., et al., Histone H4 K16Q mutation, an acetylation mimic, causes structural disorder of its N-terminal basic patch in the nucleosome. J Mol Biol, 2012. 421(1): p. 30–7.

10. Gao, M., et al., Histone H3 and H4 N-terminal tails in nucleosome arrays at cellular concentrations probed by magic angle spinning NMR spectroscopy. J Am Chem Soc, 2013. 135(41): p. 15278–81.

11. Morrison, E.A., et al., The conformation of the histone H3 tail inhibits association of the BPTF PHD finger with the nucleosome. Elife, 2018. 7.

12. Shaytan, A.K., et al., Coupling between Histone Conformations and DNA Geometry in Nucleosomes on a Microsecond Timescale: Atomistic Insights into Nucleosome Functions. J Mol Biol, 2016. 428(1): p. 221–237.

13. Biswas, M., J. Langowski, and T.C. Bishop, Atomistic simulations of nucleosomes. Wiley Interdisciplinary Reviews: Computational Molecular Science, 2013. 3(4): p. 378–392.

14. Li, Z. and H. Kono, Distinct Roles of Histone H3 and H2A Tails in Nucleosome Stability. Sci Rep, 2016. 6: p. 31437.

15. Stutzer, A., et al., Modulations of DNA Contacts by Linker Histones and Post-translational Modifications Determine the Mobility and Modifiability of Nucleosomal H3 Tails. Mol Cell, 2016. 61(2): p. 247–59.

16. Angelov, D., et al., Preferential interaction of the core histone tail domains with linker DNA. Proc Natl Acad Sci U S A, 2001. 98(12): p. 6599–604.

17. Rhee, H.S., et al., Subnucleosomal structures and nucleosome asymmetry across a genome. Cell, 2014. 159(6): p. 1377–88.

18. Gebala, M., et al., Ion counting demonstrates a high electrostatic field generated by the nucleosome. Elife, 2019. 8.

19. Gatchalian, J., et al., Accessibility of the histone H3 tail in the nucleosome for binding of paired readers. Nat Commun, 2017. 8(1): p. 1489.

20. Shaytan, A.K., et al., Hydroxyl-radical footprinting combined with molecular modeling identifies unique features of DNA conformation and nucleosome positioning. Nucleic Acids Res, 2017. 45(16): p. 9229–9243.

21. Pich, O., et al., Somatic and Germline Mutation Periodicity Follow the Orientation of the DNA Minor Groove around Nucleosomes. Cell, 2018. 175(4): p. 1074–1087 e18.

22. Gaffney, D.J., et al., Controls of nucleosome positioning in the human genome. PLoS Genet, 2012. 8(11): p. e1003036.

23. Davey, C.A., et al., Solvent Mediated Interactions in the Structure of the Nucleosome Core Particle at 1.9Å Resolution. Journal of Molecular Biology, 2002. 319(5): p. 1097–1113.

24. Macke, T.J. and D.A. Case, Modeling unusual nucleic acid structures. Molecular Modeling of Nucleic Acids, 1998. 682: p. 379–393.

25. Li, S., W.K. Olson, and X.J. Lu, Web 3DNA 2.0 for the analysis, visualization, and modeling of 3D nucleic acid structures. Nucleic Acids Res, 2019. 47(W1): p. W26–W34.

26. Rose, P.W., et al., The RCSB protein data bank: integrative view of protein, gene and 3D structural information. Nucleic Acids Res, 2017. 45(D1): p. D271–D281.

27. Pettersen, E.F., et al., UCSF Chimera--a visualization system for exploratory research and analysis. J Comput Chem, 2004. 25(13): p. 1605–12.

28. Eswar, N., et al., Comparative protein structure modeling using Modeller. Curr Protoc Bioinformatics, 2006. Chapter 5: p. Unit–5 6.

29. Galindo-Murillo, R., et al., Assessing the Current State of Amber Force Field Modifications for DNA. J Chem Theory Comput, 2016. 12(8): p. 4114–27.

30. Huang, J., et al., CHARMM36m: an improved force field for folded and intrinsically disordered proteins. Nat Methods, 2017. 14(1): p. 71–73.

31. Maier, J.A., et al., ff14SB: Improving the Accuracy of Protein Side Chain and Backbone Parameters from ff99SB. J Chem Theory Comput, 2015. 11(8): p. 3696–713.

32. Hart, K., et al., Optimization of the CHARMM additive force field for DNA: Improved treatment of the BI/BII conformational equilibrium. J Chem Theory Comput, 2012. 8(1): p. 348–362.

33. Izadi, S., B. Aguilar, and A.V. Onufriev, Protein-Ligand Electrostatic Binding Free Energies from Explicit and Implicit Solvation. J Chem Theory Comput, 2015. 11(9): p. 4450–9.

34. Onufriev, A.V. and S. Izadi, Water models for biomolecular simulations. Wiley Interdisciplinary Reviews: Computational Molecular Science, 2018. 8(2).

35. Piana, S., et al., Water dispersion interactions strongly influence simulated structural properties of disordered protein states. J Phys Chem B, 2015. 119(16): p. 5113–23.

36. Bergonzo, C. and T.E. Cheatham, 3rd, Improved Force Field Parameters Lead to a Better Description of RNA Structure. J Chem Theory Comput, 2015. 11(9): p. 3969–72.

37. Gao, K., et al., Binding enthalpy calculations for a neutral host-guest pair yield widely divergent salt effects across water models. J Chem Theory Comput, 2015. 11(10): p. 4555–64.

38. Javanainen, M., et al., Atomistic Model for Nearly Quantitative Simulations of Langmuir Monolayers. Langmuir, 2018. 34(7): p. 2565–2572.

39. Shabane, P.S., S. Izadi, and A.V. Onufriev, General Purpose Water Model Can Improve Atomistic Simulations of Intrinsically Disordered Proteins. J Chem Theory Comput, 2019. 15(4): p. 2620–2634.

40. Li, P., L.F. Song, and K.M. Merz, Jr., Systematic Parameterization of Monovalent Ions Employing the Nonbonded Model. J Chem Theory Comput, 2015. 11(4): p. 1645–57.

41. Beglov, D. and B. Roux, Finite representation of an infinite bulk system: Solvent boundary potential for computer simulations. The Journal of Chemical Physics, 1994. 100(12): p. 9050–9063.

42. Salomon-Ferrer, R., D.A. Case, and R.C. Walker, An overview of the Amber biomolecular simulation package. Wiley Interdisciplinary Reviews: Computational Molecular Science, 2013. 3(2): p. 198–210.

43. Abraham, M.J., et al., GROMACS: High performance molecular simulations through multi-level parallelism from laptops to supercomputers. SoftwareX, 2015. 1-2: p. 19–25.

44. Humphrey, W., A. Dalke, and K. Schulten, VMD: visual molecular dynamics. J Mol Graph, 1996. 14(1): p. 33–8, 27-8.

45. Phillips, J.C., et al., Scalable molecular dynamics with NAMD. J Comput Chem, 2005. 26(16): p. 1781–802.

46. Huang, H., et al., SnapShot: histone modifications. Cell, 2014. 159(2): p. 458–458 e1.

47. Khoury, G.A., et al., Forcefield_PTM: Ab Initio Charge and AMBER Forcefield Parameters for Frequently Occurring Post-Translational Modifications. J Chem Theory Comput, 2013. 9(12): p. 5653–5674.

48. Berman, H.M., et al., The Protein Data Bank. Nucleic Acids Res, 2000. 28(1): p. 235–42.

49. UniProt, C., UniProt: a worldwide hub of protein knowledge. Nucleic Acids Res, 2019. 47(D1): p. D506–D515.

50. Ikebe, J., S. Sakuraba, and H. Kono, H3 Histone Tail Conformation within the Nucleosome and the Impact of K14 Acetylation Studied Using Enhanced Sampling Simulation. PLoS Comput Biol, 2016. 12(3): p. e1004788.

51. Chakraborty, K. and S.M. Loverde, Asymmetric breathing motions of nucleosomal DNA and the role of histone tails. J Chem Phys, 2017. 147(6): p. 065101.

52. Rohs, R., et al., The role of DNA shape in protein-DNA recognition. Nature, 2009. 461(7268): p. 1248–53.

53. Kale, S., et al., Molecular recognition of nucleosomes by binding partners. Curr Opin Struct Biol, 2019. 56: p. 164–170.

54. Nacev, B.A., et al., The expanding landscape of ‘oncohistone’ mutations in human cancers. Nature, 2019. 567(7749): p. 473–478.

55. Draizen, E.J., et al., HistoneDB 2.0: a histone database with variants--an integrated resource to explore histones and their variants. Database (Oxford), 2016. 2016.

56. El Kennani, S., et al., MS_HistoneDB, a manually curated resource for proteomic analysis of human and mouse histones. Epigenetics Chromatin, 2017. 10: p. 2.

57. Skrajna, A., et al., Comprehensive nucleosome interactome screen establishes fundamental principles of nucleosome binding. Nucleic Acids Res, 2020. 48(17): p. 9415–9432.

58. Hong, L., et al., Studies of the DNA binding properties of histone H4 amino terminus. Thermal denaturation studies reveal that acetylation markedly reduces the binding constant of the H4 “tail” to DNA. J Biol Chem, 1993. 268(1): p. 305–14.

59. Phair, R.D., et al., Global nature of dynamic protein-chromatin interactions in vivo: three-dimensional genome scanning and dynamic interaction networks of chromatin proteins. Mol Cell Biol, 2004. 24(14): p. 6393–402.

60. Weaver, T.M., E.A. Morrison, and C.A. Musselman, Reading More than Histones: The Prevalence of Nucleic Acid Binding among Reader Domains. Molecules, 2018. 23(10).

61. Pilotto, S., et al., Interplay among nucleosomal DNA, histone tails, and corepressor CoREST underlies LSD1-mediated H3 demethylation. Proc Natl Acad Sci U S A, 2015. 112(9): p. 2752–7.

62. Makowski, M.M., et al., Global profiling of protein-DNA and protein-nucleosome binding affinities using quantitative mass spectrometry. Nat Commun, 2018. 9(1): p. 1653.

